# Metformin modulates autophagy in heterozygous and CRISPR-edited *TSC2* primary fibroblasts

**DOI:** 10.64898/2026.08.11.743350

**Authors:** Guilherme Danielski Viola, Pedro Ozorio Brum, Arthur Bandeira de Melo Garcia, Mariane da Cunha Jaeger, Natália Hogetop Freire, Eduardo C. Filippi-Chiela, Guilherme Baldo, Édina Poletto, Patricia Ashton-Prolla, Clevia Rosset

## Abstract

**Background:** Tuberous Sclerosis Complex (TSC) is a genetic disorder caused by variants in TSC1 or TSC2, leading to mTORC1 hyperactivation and autophagy suppression. Although TSC tumorigenesis typically follows a “two-hit” model, the role of TSC2 haploinsufficiency in autophagy regulation remains unclear. We evaluated autophagy markers in haploinsufficient and gene-edited TSC2 primary cells and investigated the role of metformin in modulating autophagy levels.

**Methods:** Primary fibroblast cultures were obtained from one healthy individual and three from patients carrying heterozygous germline TSC2 variants: the pathogenic variants c.1008T>G and c.4375C>T.A variant of uncertain significance (VUS) c.724A>T. CRISPR/Cas9-RNP editing was used to model loss of heterozygosity (LOH) in cell pools carrying each variant. Cultures were treated with rapamycin, HBSS, metformin, bafilomycin A1, or vehicle controls, and autophagy was assessed by autolysosomes formation by flow cytometry (acridine orange) and autophagosomes immunofluorescence (LC3 and p-S6K).

**Results:** In wild-type cells, only HBSS increased autophagy-positive (acridine orange-positive) cells versus control (15.6% vs. 7.5%; p=0.003). In heterozygous pathogenic cells, rapamycin and metformin increased autophagic cells: c.1008T>G (16.2%, p=0.006; 17.6%, p=0.002) and c.4375C>T (12.5%, p=0.003; 13.3%, p=0.001), versus DMSO controls (9.2% and 7.1%, respectively). VUS c.724A>T cells, with rapamycin increasing autophagic cells (9.74% vs. 6.5%; p=0.0152). In CRISPR-edited cells, all treatments increased the number of autophagic cells compared to the heterozygous cells: c.1008T>G (rapamycin 27.1% vs. 16.7%, p<0.001; metformin 27.2% vs. 17.6%, p<0.001) and c.4375C>T (rapamycin 21.3% vs. 13.1%, p=0.0021; metformin 21.5% vs. 13.6%, p=0.0029). Editing also restored metformin responsiveness in VUS cells (12.5% vs. 8.4%; p=0.0055). Immunochemistry confirmed increased total LC3II and decreased p-S6K across treated cells compared to the control (DMSO).

**Conclusion:** These findings demonstrate that TSC2 haploinsufficiency functionally impairs autophagy prior to second-hit loss. Metformin effectively restores autophagy with phenotypical changes of mTORC1 blockade, highlighting an accessible translational strategy to restore and induce autophagy in TSC cells.

## 1. Introduction

Tuberous sclerosis complex (TSC) is an autosomal dominant disorder with an estimated incidence of 1:6,000 to 1:10,000 live births [1]. The condition is defined by the widespread development of benign neoplasms, known as hamartomas, which can occur in multiple organ systems including the brain, skin, heart, lungs, and kidneys [2]. Kidney hamartomas may progress to malignancy, particularly renal cancer [3]. TSC molecular diagnosis is confirmed by presence of pathogenic variants in either the TSC1 or TSC2 genes. These genes encode the tumor-suppressor proteins hamartin and tuberin, respectively, which form a functional complex that negatively regulates the mechanistic target of rapamycin complex 1 (mTORC1), a master regulator of cell proliferation and metabolism [4, 5].

A critical downstream consequence of mTORC1 hyperactivation caused by TSC loss-of-function is the suppression of autophagy [5]. While mTORC1 physiologically suppresses autophagosome formation via ULK1 inhibition [6], the TSC loss renders this suppression constitutive, causing an autophagic block and disrupted proteostasis [7]. The role of autophagy in TSC pathogenesis is controversial. While certain TSC-associated tumors appear to be autophagy-dependent [7], impaired autophagy can promote tumorigenesis by triggering the accumulation of damaged cells [8]. Thus, the fine-tuning of autophagy in TSC appears to be one crucial process for mitigating disease manifestations.

TSC manifests in heterozygous individuals as an autosomal dominant disease. However, previous studies suggest that TSC tumorigenesis follows the classical Knudson ‘two-hit’ model, which requires somatic loss of the remaining functional allele (loss of heterozygosity, LOH) for tumorigenesis [9]. Conversely, the haploinsufficient state is sufficient to trigger autophagy dysregulation in TSC cells [8] and mTORC1 hyperactivation [10]. Therefore other cellular cellular pathways in addition to mTORC1 are altered in the haploinsufficient state, even before LOH occurrence, and these pathways could be targeted to modulate disease progression.

Rapamycin and its analogs are FDA-approved for TSC by directly inhibiting mTORC1. However, optimal duration of treatment and long-term dosing requirements remain unknown, but complete interruption of mTORC1 inhibitor therapy for TSC-related tumors typically results in lesion regrowth [11]. Moreover, chronic mTORC1 inhibition induces adaptive resistance, most notably through the release of the negative feedback loop on insulin receptor substrate 1 (IRS-1), culminating in a paradoxical, compensatory activation of the pro-survival PI3K/AKT pathway mediated by mTORC2 [12]. Therefore, targeting upstream and downstream pathways presents an alternative approach in TSC treatment. In this study, we aimed to evaluate autophagy pathway status across distinct *TSC2* genetic contexts, such as heterozygous cells harboring pathogenic variants or a variant of uncertain significance (VUS), and CRISPR/Cas9-edited cells modeling loss of heterozygosity (LOH), in response to different autophagy modulators, including the mTORC1 inhibitor rapamycin, nutrient starvation (HBSS), and metformin. Given the close connection between autophagy and mTORC1 signaling, and its potential as a therapeutic target in TSC, we sought to determine whether TSC2 haploinsufficiency alone is sufficient to dysregulate autophagy, and how this response is further shaped by LOH and by metformin treatment.

## 2. Materials and Methods

### 2.1 Skin biopsy and molecular diagnosis

Biopsies (6 mm) of normal-appearing skin were obtained from three patients diagnosed with TSC according to clinical criteria [1] and one healthy individual. The molecular diagnosis strategy previously used to identify germline variants in TSC2 was a customized next generation sequencing panel followed by multiplex ligation probe-dependent amplification (MLPA) [8]. All patients were included in this study after an interview and the signing of a specific written informed consent form. The study was approved by the Research Ethics Committee of the Hospital de Clínicas de Porto Alegre (CEP-HCPA) under protocol number CAAE 10508919.7.0000.5327.

Three patients were confirmed with clinical and molecular diagnosis criteria of Tuberous Sclerosis Complex (TSC) in the TSC2 gene (NM_000548.5, ENST00000219476.9) [1]. Two of these patients carried TSC2 nonsense variants previously classified as pathogenic (c.1008T>G and c.4375C>T), while the third harbored a variant of uncertain significance (VUS) (c.724A>T).

Primary fibroblasts derived from an unaffected family member served as the wild-type control. More information about the patients is present in Table 1.

**table 1.**
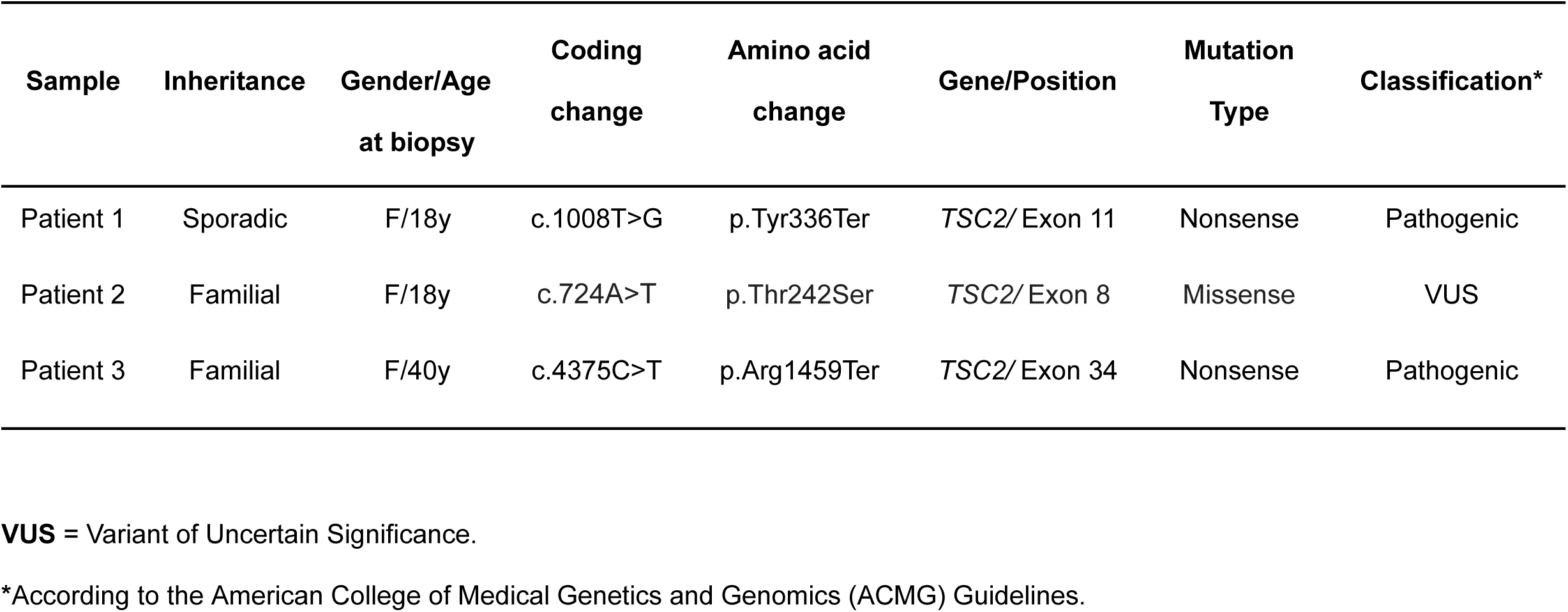
Variant analysis of TSC patients included in this study.

### 2.2 Cell culture and treatment

Primary fibroblasts isolated as aforementioned were grown in HAM-F10 medium (ThermoFisher Scientific, USA) supplemented with 1% penicillin-streptomycin (ThermoFisher Scientific, USA) and 20% fetal bovine serum (Gibco Laboratories, USA), in 25cm² cell culture flasks incubated at 37°C in 5% CO2 and 95% air humidified atmosphere. When cells ((up to the 7th passage) appropriate confluence (80-90%) the cell culture media was removed and cells were treated with 1.5 mL trypsin (0.25%) (Gibco Laboratories, USA) for 5 minutes at 37°C. Supplemented medium (1.5 mL) was added to block trypsin action, and the detached cells were collected in a 15 mL Falcon tube and centrifuged at 500g for 5 minutes. The supernatant was discarded, and pellet cells were suspended in 1 mL medium and plated at appropriate number/confluency for experiments.

After confluence, cells were treated with rapamycin 100nM (R0395 - Sigma Aldrich, USA) 10 mM metformin (PHR1084 - Sigma Aldrich, USA), Hanks Balanced Salt Solution (HBSS) (Gibco, USA) and vehicle (DMSO) for 12 hours and 100 nM bafilomycin A1 (B1793 - Sigma Aldrich, USA) for 2 hours.

### 2.3 RNP-CRISPR/Cas9-RNP editing

A gene knock-out kit containing three sgRNAs (UCUUUAGGGCGAGCGUUUGG/CGUGAAGGUCUUCGUUGGAA/UGGGAGACACAUC ACCUACU; (GKOv2) from Synthego (Synthego Corporation, USA) was designed to target the TSC2 gene. For ribonucleoprotein (RNP) complex formation, 1 μl of Cas9 20μM (Synthego Corporation, USA) was incubated with 6 μl of gRNA (30μM) (9:1 gRNA:Cas9 ratio). After incubation, 7ul of the RNP complex was combined with 1.5×10⁵ cells resuspended in 18ul of 1M buffer in microcentrifuge tubes. For electroporation control, 1.5×10⁵ of the primary cells were resuspended in 24,6 μl of 1M buffer and 0,4 μg of the reporter plasmid (pmaxGFP vector). Nucleofection was performed in the 4D Lonza Nucleofector X Unit using the EH100 program protocol.

### 2.4 Genome editing confirmation

Total isolated DNA was from cultured transfected cells using QIAamp DNA Mini Kit (QIAGEN). The total DNA obtained from cell cultures were amplified in a 20 μL volume PCR reaction: 0.5µl Forward primer (10nM)(AATGCTGATGCTGCAGACCT), 0.5µl Reverse primer (10nM) (ATGAAGCAGGGTGGGCATAC), ultrapure water, 2.5µl buffer with Tris-HCl 200 mM, pH 8,4, KCl - 500 mM without MgCl2 (Invitrogen), 0.5-0.8µl dNTP, 0.5-0.8µl MgCl2, 0.1µl Platinum Taq DNA polymerase (Invitrogen™), and 1µl of DNA. PCR reaction was performed in Proflex thermal cycler using the following parameters: denaturation at 94°C for five minutes, followed by 35 cycles at 94°C for 30 seconds, 60°C for 45 seconds, 72°C for 30 seconds, and a final extension step of 72°C for two minutes. For correct TSC2 amplicon verification, electrophoresis step was performed in 1% Agarose Gel (Uniscience Corporation) embedded in TBE 1X buffer. PCR amplicons were purified in a reaction containing 5μL of amplicons and 2μL of ExoSAP-IT (ThermoFisher Scientific, USA) for 15 minutes at 15°C and 30 minutes at 80°C in a Proflex thermal cycler.

Purified PCR products were submitted to Sanger sequencing. PCR products were labeled with 5.0 pmol of the primer 5’-AATGCTGATGCTGCAGACCT-3’ and and 1 uL of BigDye Terminator v3.1 Cycle Sequencing Kit (Applied Biosystems, CA, United States) in a final volume of 10 uL. Labeling reactions were performed in a Veriti 96-Well Thermal Cycler (Applied Biosystems, CA, United States) thermocycler with an initial denaturation step of 96°C for 1 min followed by 35 cycles of 96°C for 15 sec, 50°C for 15 sec and 60°C for 4 min. Labeled samples were purified using BigDye XTerminator Purification Kit (Applied Biosystems) and electron injected in the genetic analyzer ABI 3500 Genetic Analyzer (Applied Biosystems, CA, United States) with 50 cm capillaries and POP7 polymer (Applied Biosystems, CA, United States). Sanger electropherogram results in .ab1 format were checked for FinchTV 1.4 (https://digitalworldbiology.com/FinchTV).

The Synthego ICE Analysis tool (v3) (Synthego Performance Analysis, ICE Analysis, https://www.synthego.com/products/bioinformatics/crispr-analysis) was performed to estimate genome editing frequency. Sanger .ab1 format results and information about gRNAs were uploaded into the ICE software. Assessment of genome editing was performed by comparing the frequency of indels of the unedited vs edited cells.

### 2.5 Flow cytometry

Each primary fibroblast cell was seeded as 5×10⁴ per well in triplicates in a 24-well plate. Cells were centrifuged for 5min at 500g and washed with PBS 1x three times. Subsequently, cells were resuspended in 500 μL HAM-F10 medium and incubated for 15 minutes with acridine orange (AO) solution (1 μg/mL). The percentage of autophagic cells was measured in Attune NxT flow cytometer with BL3-A vs BL1-A axis (ThermoFisher Scientific, USA).

### 2.6 Immunofluorescence

Cells were seeded at 5×10⁴ in a 24-well plate. After the treatment, cells were washed with 500 µl of 1X PBS three times and then fixed in 4% paraformaldehyde for 10 min at room temperature, followed by 10-min permeabilization utilizing ice-cold 0.1% Triton-PBS. To block nonspecific binding, cells were incubated with 3% albumin (BSA), in T-PBS (PBS + Tween 20 0.1%) for 1 h at room temperature. Cells were then incubated overnight with primary antibodies with Anti-LC3B antibody ab63817 (1:400) and anti-p-p70 S6 kinase a (A-6): sc-8416 (1:200), followed by a one hour incubation with its species-specific corresponding secondary antibody coupled with Alexa Fluor® staining (488 nm) from Cell Signalling Technology at room temperature (all secondary antibodies were used at 1:1000 dilution).

Images were obtained using a fluorescence microscope Olympus BX51. For each condition (treatment and replicate) at least one picture (field) of each quarter of the coverslip was taken, representing the entire coverslip confluency and immunolabelling, accounting no less than one hundred viable cells for each slide in each replicate. The pictures used for analysis were taken at the same light exposition with a 20× objective (200× magnification).

Whole-cell and nuclear masks were generated in Fiji using color-based thresholding, with threshold parameters manually adjusted for each image. Nuclear segmentation was based on the DAPI channel, while whole-cell segmentation was performed using a composite of the remaining fluorescence channels. Individual cell ROIs were extracted from the resulting binary masks using the Analyze Particles function. Cytoplasmic ROIs were derived by subtracting the nuclear mask from the whole-cell mask. Integrated density was used as a measure of fluorescence intensity for both nuclear and cytoplasmic compartments to ensure comparability across regions. 3-5 images were quantified per biological replicate.

### 2.7 Statistical analysis

Statistical analysis was performed with the GraphPad Prism software version 10.0 (GraphPad Software Inc., San Diego, USA). Data was evaluated by unpaired Student’s t-test or One-way ANOVA followed by Tuckey test when suitable. Differences were considered significant when p < 0.05.

## 3. Results

### 3.1 TSC2 heterozygous primary fibroblasts expresses changes in autolysosome amount

We examined the autolysosomes formation in order to screen the proportion of autophagic cells in TSC2 heterozygous and wild-type conditions following treatment with autophagy-inducing compounds. In wild-type fibroblasts, the analysis of autolysosomes by acridine orange (AO) staining followed by flow cytometry revealed that only HBSS significantly increased the proportion of AO-positive cells compared to the DMSO control (15.6% vs. 7.5%; p=0.003) (Figure 1a).

**Figure 1:**
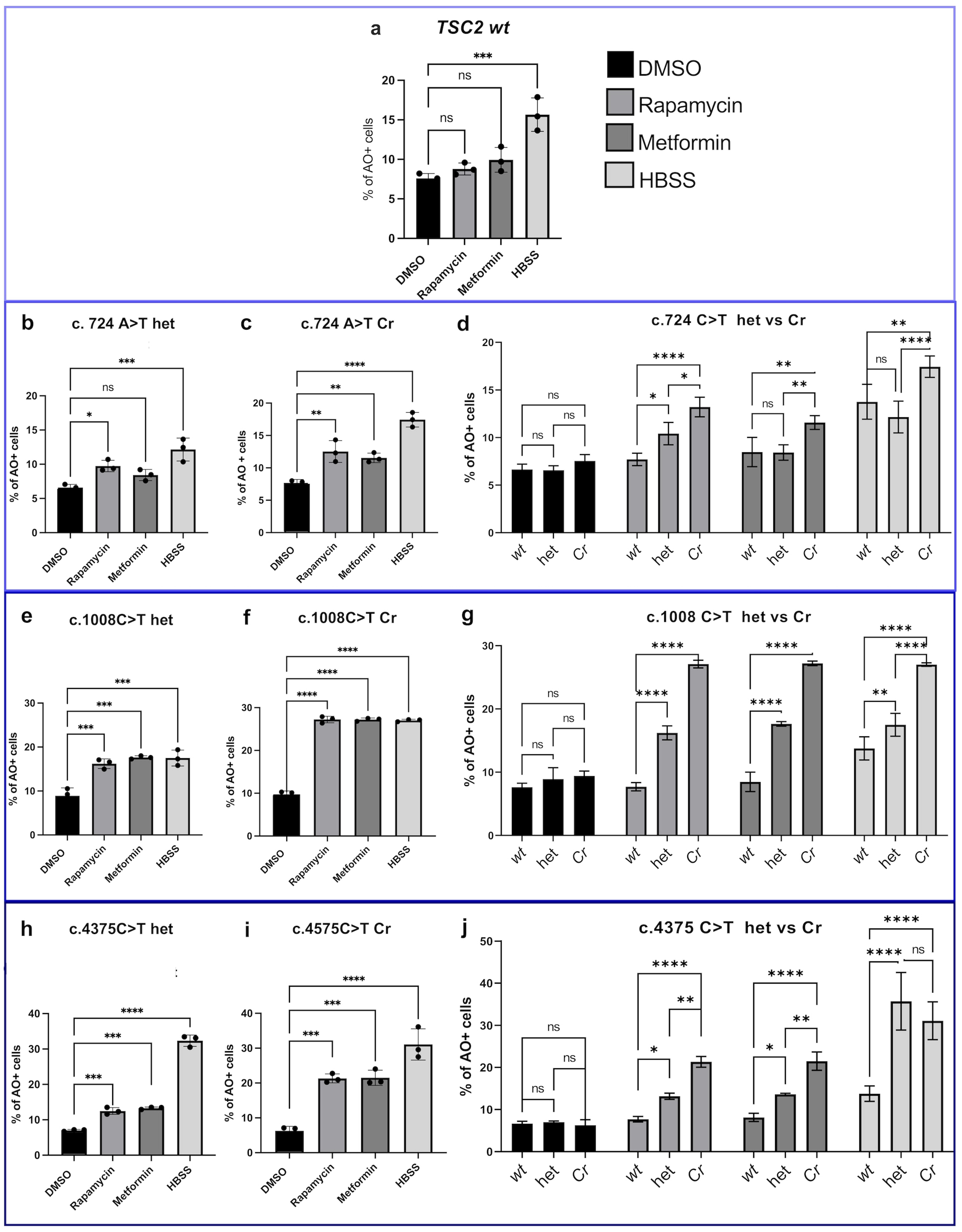
Proportion of autophagic cells measured by flow cytometry with acridine orange (AO). Columns represent mean ± SD. Each dot represents the mean percentage of autophagic cells in the biological replicate. het*=TSC2* variant present; Cr = het cells that underwent *TSC2* CRSPR gene editing. (one-way ANOVA followed by Tukey’s Multiple Comparison post-hoc test *p < 0.05, **p < 0.01, ***p < 0.001, **** p < 0.0001, ns = not significant)

Regarding the heterozygous TSC2 c.724A>T cells, rapamycin (9.74%; p=0.0152) and HBSS (12.15%; p=0.0005) increased the proportion of autophagic cells compared to DMSO (6.5%)(Figure 1b). For the c.1008T>G variant, all treatments increase in autophagic cells compared to DMSO (9.2%): rapamycin (16.2%; p=0.006), metformin (17.6%; p=0.002), and HBSS (17.5%; p=0.002)(Figure 1e). Similarly, in c.4375C>T cells, all treatments increased OA-positive cells proportion compared to the DMSO group (7.1%). Specifically, the proportion of autophagic cells was 12.5% with rapamycin (p=0.003), 13.3% with metformin (p=0.001), and 32.4% with HBSS (p<0.0001) (Figure 1h).

### 3.2 CRISPR/Cas9-mediated TSC2 editing enhances autolysosome formation

After confirming the sensitivity to autophagy induction conferred by the TSC2 heterozygous variant in the primary cells, we hypothesized that the loss-of-function of the second allele (LOH) could enhance the treatment-induced autophagy. Cells underwent CRISPR/Cas9-RNP genome editing with three single guide RNA for higher editing accuracy. After edition, the percentage of indels in the mixture of the primary cells edited-pool acquired were c.724A>T, 17% of edited cells (50% of the edited pool with fragment deletion of 150pb and 80pb); c.1008T>G, 27% (22% of the edited pool with - 1 deletion), c.4375C>T 14% (70% of 14% was a fragment deletion of 156pb) (see more edited population in sup figure 1).

Upon editing, c.724A>T cells exhibited a higher percentage of autophagic cells compared to the heterozygous state under identical treatment conditions. Metformin, which demonstrated no response in the heterozygous state, induced a significant increase in autophagy in edited cells (12.5% vs 8.4%; p=0.0055) (Figure 1c). Specifically, rapamycin increased the autophagy proportion from 10.4% in heterozygous cells to 13.2% in edited cells (p=0.0138) (Figure 1d). HBSS treatment resulted in 17.4% vs. 12.1% (p<0.0001) in heterozygous versus edited cells, respectively (Figure 1d). Notably, DMSO treatment showed no significant difference between the heterozygous vs edit groups (p=0.52) (Figure 1d).

For the c.1008T>G variant, all treatments in increased autophagy levels in edited cells compared to heterozygous: rapamycin (27.1% vs 16.7%; p<0.001), metformin (27.2% vs 17.6% ; p<0.001), and HBSS (27.0% vs 17.5%; p<0.001) (Figure 1g. Results for the c.4375C>T variant followed a similar trend for rapamycin (21.32% vs 13.1%; p=0.0021) and metformin (21.5% vs 13.6%; p=0.0029). However, HBSS treatment showed no statistically significant difference (35.7% vs. 31.1%; p=0.09)(Figure 1j).

### 3.3 Morphological and quantitative analysis of autophagosomes

The assessment of the LC3-II marker in autophagosome membranes through immunochemistry is a standard method for monitoring autophagy [14]. We evaluated autophagosome formation through LC3-II visualization in addition to autolysosome analysis as complementary methods to evaluate autophagy, allowing the screening of the initial and final step of this process, respectively. To analyze LC3-II in autophagosome membranes, we subjected both heterozygous and TSC2-edited cells to the same conditions (DMSO, rapamycin, metformin, and HBSS) in addition to bafilomycin, and rapamycin + bafilomycin to enable autophagic flux evaluation. LC3-II was visualized via green fluorescence, allowing for a qualitative assessment of its intensity across treatments (Figure 2). Considering the dynamic nature of autophagy, relevant information comes from the distinction between altered autophagosome biogenesis or impaired autophagic flux. The use of autophagy markers such as LC3-II can be complemented by analysing the overall autophagic flux to permit a correct interpretation of the results. To evaluate autophagy flux, we used bafilomycin A1, which causes an increase in lysosomal/vacuolar pH, and, ultimately, blocks fusion of autophagosomes with the vacuole. Phenotypic heterogeneity was observed regarding cell size, morphology, and fluorescence signal intensity. Furthermore, the basal autophagy was evident in the micrographs, characterized by a variable pattern of LC3-II staining in individual cells.

**Figure 2:**
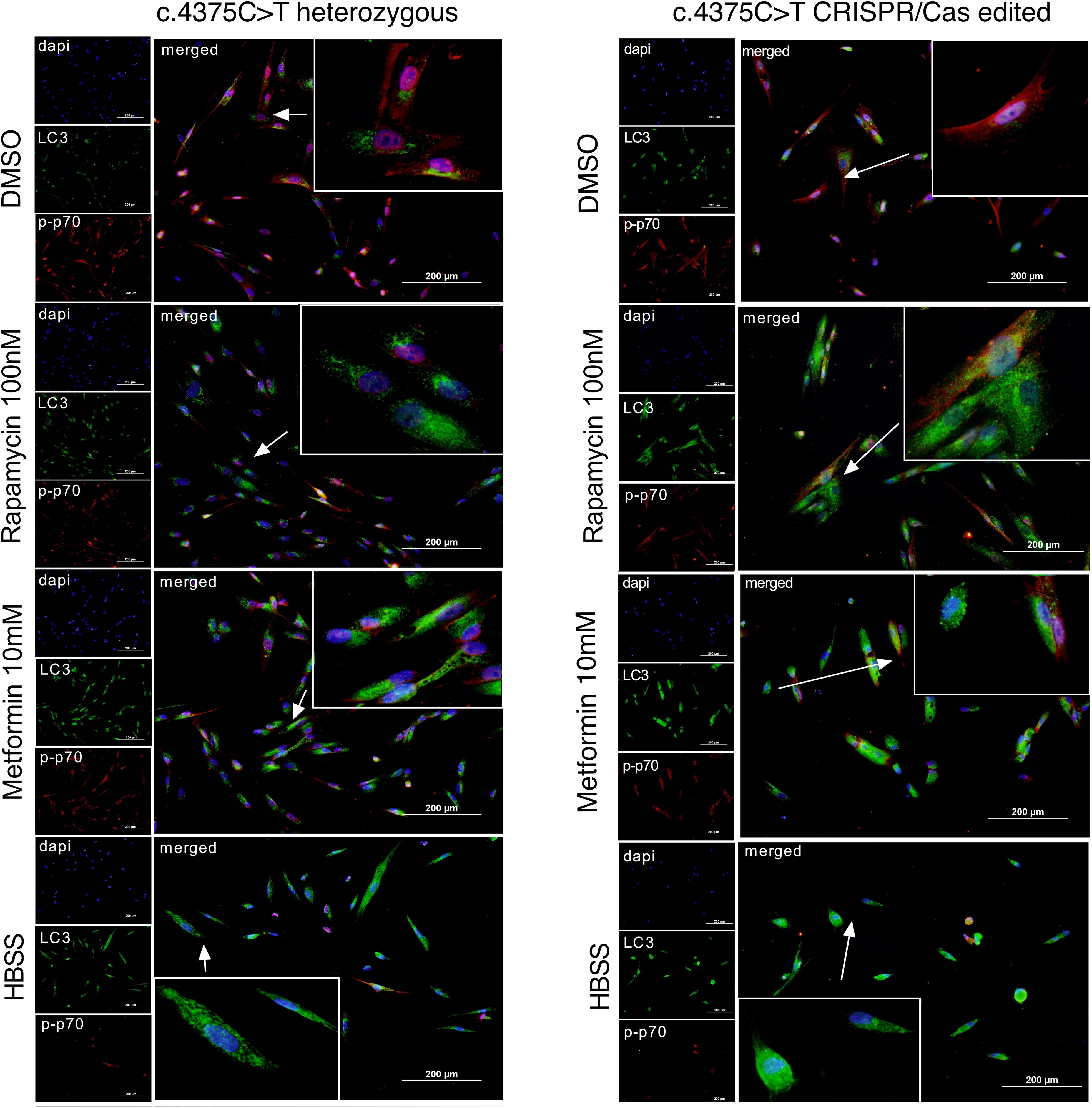
Autophagosome imaging. Representative immunoflorescence images of the *TSC2* c.4375 C>T in heterozygous state and CRISPR edited. Dapi is shown in blue, p-p70 (p-S6K) in red and LC3 in green fluorescence. The arrows indicate the zoomed cell/cells for autophagosomes emphasis. All photos were taken in 20X objective and the zoom at 40x objective.

Visual inspection revealed that the DMSO control did not indicate morphological differences between heterozygous and gene-edited cells. In contrast, rapamycin appeared to increase the number and formation of larger autophagosomes (Figure 2). Conversely, metformin and HBSS treatments caused a reduction in cell width across all cell lines compared to the DMSO control. This phenotypic alteration was accompanied by a higher frequency of autophagosomes that were, however, smaller in size compared to those induced by rapamycin. Bafilomycin, used as an autophagic flux control to inhibit autophagosome-lysosome fusion, led to an accumulation of enlarged autophagosomes without significantly increasing their total number or the proportion of positive cells, confirming functional autophagic flux (see sup Figure 2). Regarding p-S6K, a downstream protein of mTORC1 in the mTOR signalling pathway and a marker of ribosome activity, showed a reduced signal intensity in groups treated with autophagy-inducing compounds (rapamycin, metformin, and HBSS) compared to DMSO (Figure 2).

Complementary data provided by flow cytometry, such as cell size (FSC-H), demonstrated changes in this surrogate marker (decreased size) under all treatment conditions (rapamycin, metformin, and HBSS), consistent with the phenotypic and morphological changes observed in both heterozygous and CRISPR-edited cells (data not shown)

Interestingly, for VUS patient 1, primary cells harboring the c.724A>T variant demonstrated the same pattern for the treatments when comparing the green fluorescence from heterozygous cells and the CRISPR-edited pool. Only the DMSO vehicle control showed no alterations in LC3-II fluorescence (p=0.9904)(Figure 3a). In addition, when comparing the p-S6K fluorescence among treated groups, the edited pool revealed a decreased protein signal in cells treated with rapamycin or metformin (p= <0.0001; p= <0.0001) (Figure 3b).

**Figure 3.**
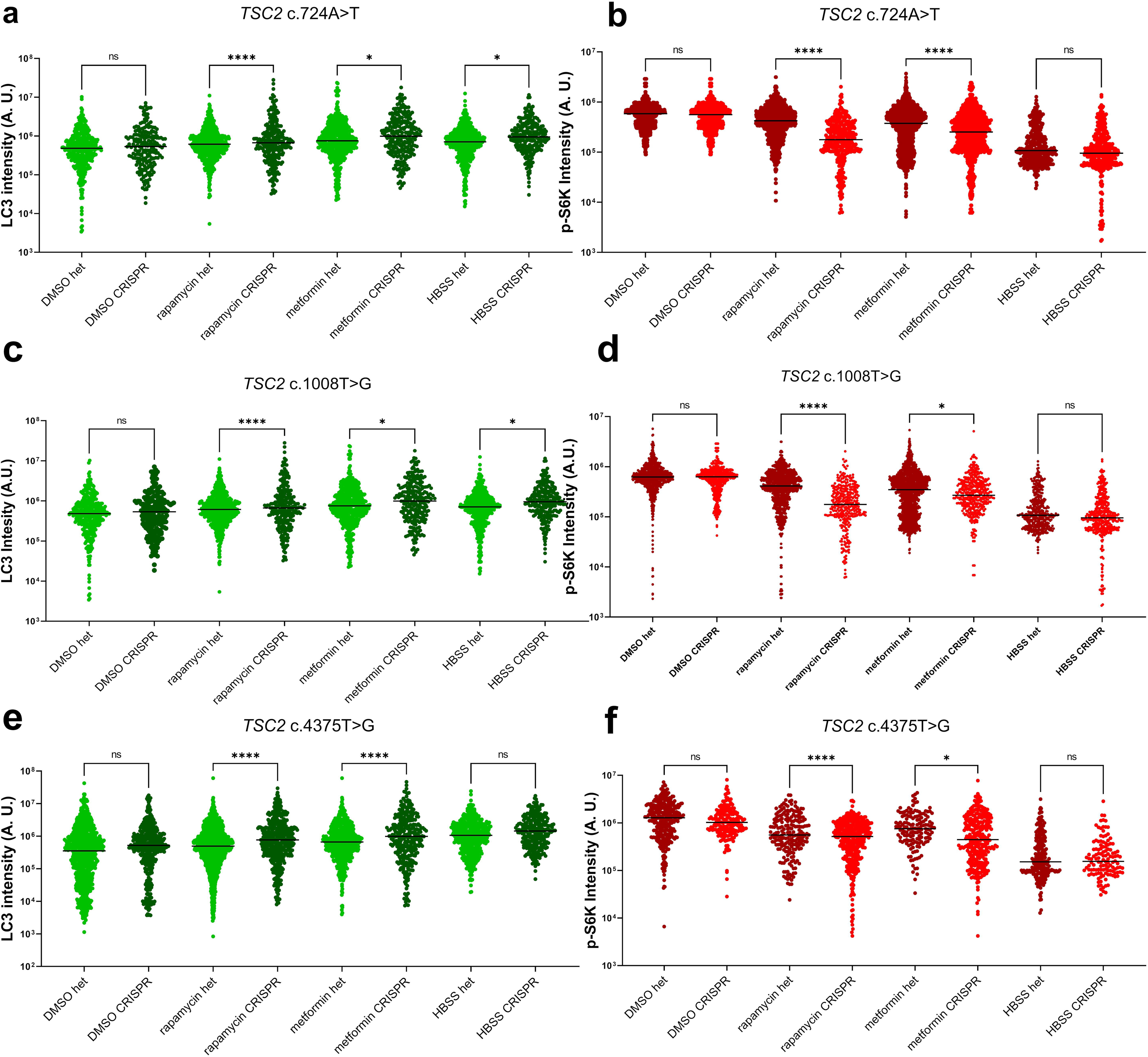
LC3II and p-S6K cellular content comparison between treatments. Immunofluorescence quantification analysis. a,c,e represents the LC3 (green) and b,d,f p-S6K (red) intensity quantified (raw intensity) and plotted as arbitrary units (A. U.). Each dot represents the fluorescence intensity of one individual cell one-way. Multiple t-tests were used for statistical analysis comparing the heterozygous vs CRISPR group for each treatment. Results were considered significant when p<0.05 (*p < 0.05, **p < 0.01, ***p < 0.001, **** p < 0.0001)

The quantification of the raw integrated density of cytoplasmic LC3-II in c.1008T>G cells showed no significant baseline differences between heterozygous and edited cells in the DMSO group (p=0.7823). However, the edited cells exhibited elevated LC3-II protein levels across rapamycin, metformin, and HBSS (Figure 3c). Furthermore, the quantification of nuclear p-S6K revealed a decreased protein signal in cells treated with rapamycin, metformin (p=<0.0001; 0.0269) (Figure 3d).

However, for the pathogenic c.4375T>G variant, DMSO and HBSS demonstrated no alteration among the heterozygous vs the CRISPR edited pool (p=0.7823, p=0.0561, respectively) when once again the rapamycin, metformin treatment increased the cytoplasmic intensity fluorescence of LC3-II (p<0.0001, p<0.000.1)(Figure 3e). While LC3-II increased in these groups, the p-S6K fluorescence demonstrated the opposite scenario with diminished nuclear fluorescence in the rapamycin, metformin groups (p<0.0001, p<0.0001, Figure 3f) no differ

For all the tested cells with different TSC2 variants, bafilomycin A1 treatment significantly decreased LC3-II levels compared to rapamycin (Figure 4).The treatment with rapamycin plus bafilomycin A1 increased total LC3-II levels (Figure 4) compared to bafilomycin A1 alone, suggesting that the treatment may increase the synthesis of autophagy-related membranes (which seems to be impaired in TSC cells) instead of provoking alterations in autophagy flux.

**Figure 4.**
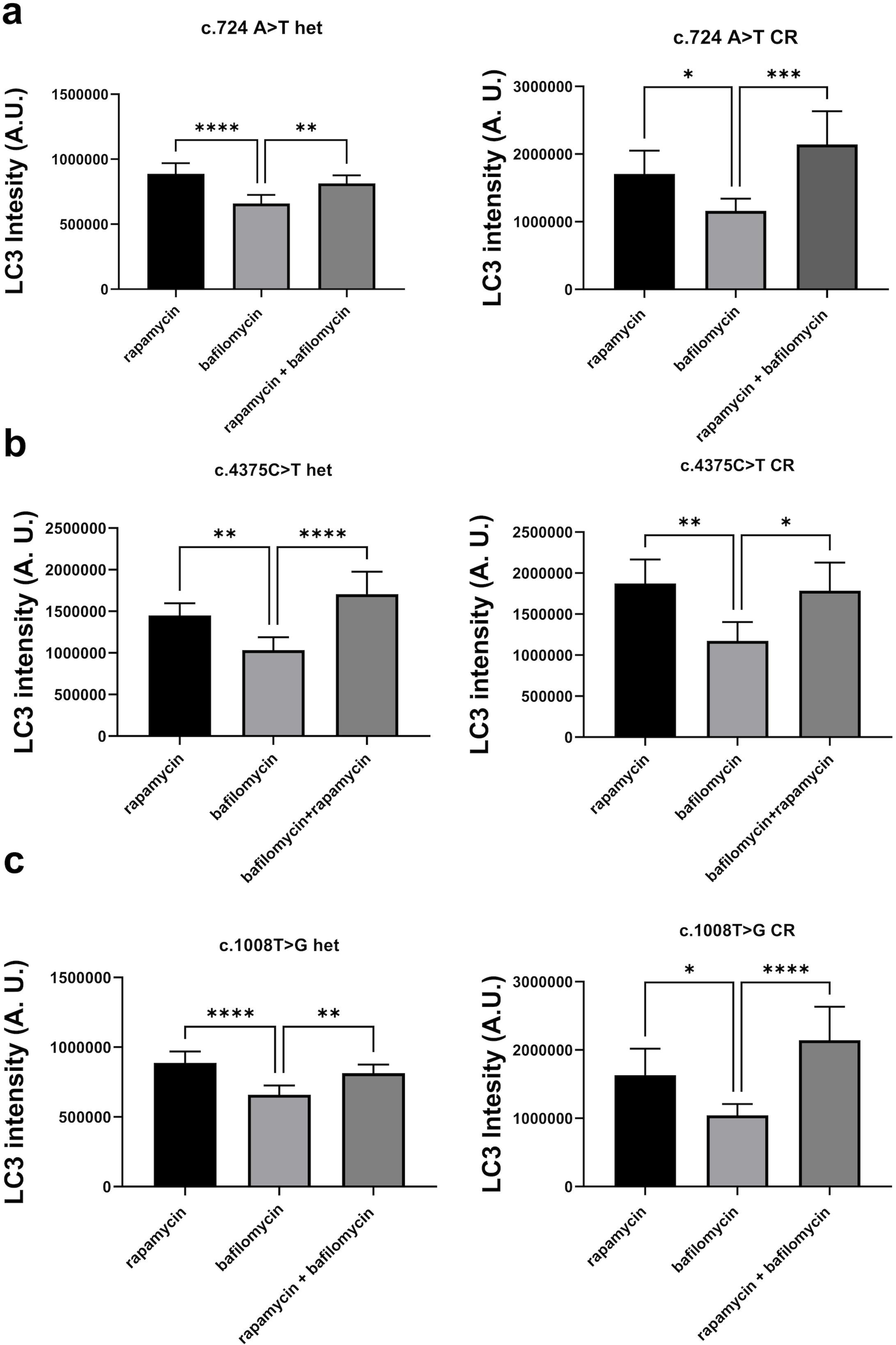
Assessment of autophagic flux in primary fibroblasts harboring TSC2 variants. LC3 fluorescence intensity (expressed in Arbitrary Units - A.U.) was quantified to determine autophagic flux dynamics across three distinct variants: (a) c.724 A>T; (b) c.4375C>T; and (c) c.1008T>G. For each variant, experiments were performed in heterozygous (het) and CRISPR-edited (CR) cells. Cells were treated with Rapamycin, Bafilomycin A1, or the combination of both compounds. Data represent mean ± SEM of intensity from each individual cell quantified. Statistical analyses were performed using One-way ANOVA, with significance indicated as: * p < 0.05; ** p < 0.01; *** p < 0.001; **** p < 0.0001.

## 4. Discussion

Understanding the molecular mechanisms involved in TSC pathogenesis remains a challenge due to its phenotypic heterogeneity regarding tumor location, size, and type, as well as the complexity of the mTORC1 and its related pathways. Although loss-of-function of the second allele in tumor suppressor genes explains tumor formation in many hereditary cancer syndromes, evidence suggests that tumors can occur in the presence of TSC1/TSC2 variants in a heterozygous state, as demonstrated in subsets of TSC-associated lesions lacking an identifiable second hit [14]. This reinforces the role of haploinsufficiency in the disease scenario. Importantly, the cellular processes that are connected and altered due to mTORC1 hyperactivation in the haploinsufficient state could be modulated in conjunction with mTORC1 inhibition.

In this work, the use of TSC2 wild-type cells played a critical role as a control for basal autophagy levels. When treated with rapamycin or metformin alone, wild-type cells did not exhibit significant alterations in autophagy levels, as measured by autolysosome formation. This result was expected, as normal mTOR signaling maintains physiological basal autophagy levels in non-hyperactive cells. However, exposure to HBSS for 12 hours robustly induced autophagy in wild-type cells, as evidenced by increased autolysosome formation and marked changes in cell cytomorphology. Notably, HBSS stimulation induced a cellular phenotype characterized by nutrient deprivation stress in wild-type cells. It is important to mention that in addition to wild-type TSC cells used as control, we used a vehicle control treatment (DMSO), which showed no significant variation in autophagy levels across all cell lines tested, validating the specificity of compound-induced autophagy modulation.

Our findings demonstrated that not only do TSC2 wild-type cells exhibit different autophagy sensitivity to stimuli compared to TSC2 heterozygous variant-harboring cells, but the type of variant also generates variability in the autophagic outcome. Cells harboring known pathogenic nonsense variants in critical functional domains, such as exon 11 (the hamartin-binding domain) and exon 34 (the GAP domain), exhibited high autophagic induction upon stimulus. Furthermore, a novel and significant observation was the intermediate phenotype elicited by the c.724A>T variant, a missense alteration in the exon 8 (hamartin interaction region), a variant of uncertain significance (VUS). Heterozygous cells with this variant displayed a significant increase in autophagy compared to wild-type cells when rapamycin treated, but no alteration when metformin treated.

The results obtained in TSC2 gene-edited cells suggest that LOH enhances autophagy dysregulation in the context of TSC disease. The TSC2 biallelic loss-of-function triggers a stronger response to rapamycin and metformin treatments when compared to heterozygous TSC2 cells, indicated by a higher increase in the proportion of autophagic cells in LOH cells after treatment. This finding reveals that autophagy basal levels are dysregulated in heterozygous cells and this dysregulation is amplified by LOH. Direct mTORC1 inhibition with chronic rapamycin exposure in certain primary fibroblasts can induce an hypertrophic phenotype linked to cellular senescence or massive autophagosome accumulation [16].

Furthermore, p-S6K immunoassay results correlated to the LC3-II in a logical and negative way, as expected. In the cells with reduced autophagy levels (variant-harboring cells before treatments), the levels of p-S6K were increased. In cells where the autophagy levels were enhanced after autophagic stimulating compound treatment, the levels of p-S6K were reduced. Moreover, morphological characteristics such as decreased volume/size together with increased number of observed autophagosomes contribute to the effects observed by the treatment and a distinct phenotype.

Altogether, our results based on autophagy modulation suggest that metformin could be used to modulate this process in TSC heterozygous and edited. While rapamycin acts as a direct allosteric mTORC1 inhibitor, metformin, a widely used anti-diabetic drug, activates AMP-activated protein kinase (AMPK) and indirectly inhibits mTORC1 signaling by mimicking a state of cellular energy depletion [17, 18]. In fact, recent clinical data showed that metformin did significantly reduce SEGA volume and seizure frequency compared with placebo [19]. However, none of the previous studies in TSC models assessed the precise mechanism by which metformin acts in TSC control and none investigated autophagy after metformin treatment. We hypothesize that metformin appeared to act via bioenergetic stress pathways, such as AMPK-driven autophagy. This distinction aligns with known molecular networks where the AMPK-mTOR pathway regulates cell fate determination through TFEB-dependent autophagic regulation [20]. Hence, metformin alters autophagy levels in a TSC-deficient background, but may do so through a metabolic stress response that is phenotypically and mechanistically distinct from direct mTORC1 inhibition. This distinction is particularly relevant given rapamycin’s clinically observed side effects, such as mucositis and metabolic dysregulation, which limit its long-term tolerability and efficacy.

In this study, only the cells harboring a TSC2 VUS did not respond to metformin in the same way as the cells harboring a pathogenic variant. This suggests that this variant, in a heterozygous state, may have basal autophagy levels near to the wild-type cells in a nutrient-rich environment. However, after edition, the LOH VUS cells responded to the metformin treatment, suggesting that variants with less effect on TSC protein function could be more dependent on the second hit to trigger autophagy dysfunction and also disease complications. In this line, these results provide functional in vitro insights that could be used in conjunction with the American College of Medical Genetics and Genomics (ACMG) criteria [21] for the clinical interpretation of VUS in TSC.

This study has limitations and methodological considerations. Extensive data demonstrate the challenges of gene editing in primary cells. Unlike immortalized cell lines, primary cells are highly susceptible to Cas9-induced double-strand breaks, which frequently trigger a p53-mediated DNA damage response leading to cell cycle arrest or apoptosis [22, 23]. Furthermore, human primary fibroblasts are intrinsically prone to stress-induced premature senescence (SIPS) when subjected to standard in vitro culture conditions and the substantial physiological stress of electroporation [24, 25]. Given these intrinsic biological limitations, we established a maximum of one month of cell culture after transfection. In this period, the editing efficiency achieved in our primary cell pool was 14-27% (up to 27% of cells carried LOH in a pool). Importantly, this heterogeneous cell pool containing both TSC2 heterozygous and LOH cells actually reflects the disease scenario, where not all cells in a lesion harbor LOH. The primary cell culture limitation did not allow additional cell replications and the seeding of a specific proportion of LOH cells in each well. However, this approach enabled us to capture autophagy dysregulation across different disease states while maintaining cellular integrity.

Another methodology limitation is relative to the LC3-II evaluation. Our approach used immunofluorescence visualization of LC3-II on autophagosomal membranes as a reliable form to assess autophagosome formation and dynamics without requiring protein extraction and densitometric analysis. In addition, we performed an additional flow cytometry with acridine staining, which is complementary to the immunofluorescence and provides morphometrical characteristics of the treated cells. Acridine orange, a cationic dye, selectively accumulates in acidic compartments (autolysosomes), while immunofluorescence detection of LC3-II localization on autophagosomal membranes offers superior spatial resolution and morphological information compared to blotting techniques alone.

Our strategy provided important functional in vitro evidence that contributes to the understanding of TSC pathogenesis and supports the previous and further clinical investigation of metformin as an alternative or adjunctive therapeutic approach for TSC management. The findings suggest that metformin may function as a preventive therapeutic modulator in the germline context of TSC, where all cells constitutively carry TSC2 heterozygous pathogenic variants. Functional characterization of additional variants across the TSC2 gene would strengthen our conclusions regarding variant-dependent autophagy phenotypes. Moreover, our findings are restricted to autophagy-related processes, and complementary studies investigating additional pathways directly or indirectly related to mTORC1 would provide a more comprehensive understanding of metformin therapeutic potential in TSC.

## 5. Conclusion

In conclusion, our findings demonstrate that basal autophagy levels are dysregulated in heterozygous cells carrying pathogenic *TSC2* variants alongside mTORC1 hyperactivation, an effect further amplified by loss of heterozygosity (LOH). Although this autophagic imbalance is less pronounced in heterozygous VUS cells, edited cells trigger a significant functional defect. Notably, metformin modulates autophagy dysregulation in both haploinsufficient and LOH states. By operating through mechanisms distinct from direct mTORC1 inhibition, metformin highlights its potential as an alternative or complementary therapeutic strategy in TSC, justifying further exploration of autophagy-targeted compounds.

## Acknowledgments

We gratefully thank Fundo de Incentivo à Pesquisa e Eventos (FIPE) of Hospital de Clínicas de Porto Alegre, Fundação de Apoio à Pesquisa do Rio Grande do Sul (FAPERGS) for the process 24/2551-0001231-1 and Coordenação de Aperfeiçoamento de Pessoal de Nível Superior (CAPES) for their financial support. The author Patricia Ashton-Prolla acknowledges the National Council for Scientific and Technological Development (CNPq) for the Research Productivity Fellowship 1B.

## CRediT authorship contribution statement

**Guilherme Danielski Viola:** conceptualization, investigation, formal analysis, writing – original draft. **Pedro Ozorio Brum:** conceptualization, investigation, formal analysis. **Arthur Bandeira de Melo Garcia:** conceptualization, investigation, formal analysis. **Mariane da Cunha Jaeger:** conceptualization, investigation, resources. **Natália Hogetop Freire:** investigation **Eduardo C. Filippi-Chiela:** conceptualization, writing – review & editing. **Guilherme Baldo:** conceptualization, writing – review & editing. **Edina Poletto:** conceptualization, writing – review & editing. **Patricia Ashton-Prolla:** conceptualization, formal analysis, writing – review & editing. **Clevia Rosset:** conceptualization, investigation, formal analysis, writing – review & editing.

All authors read and approved the final version of the manuscript.

## References

1. Northrup H, Aronow ME, Bebin EM, Bissler J, Darling TN, de Vries PJ, et al. Updated international tuberous sclerosis complex diagnostic criteria and surveillance and management recommendations. Pediatr Neurol. 2021;123:50–66.

2. Schwartz RA, Fernández G, Kotulska K, Jóźwiak S. Tuberous sclerosis complex: advances in diagnosis, genetics, and management. J Am Acad Dermatol. 2007;57(2):189–202.

3. Henske EP, Cornejo KM, Wu CL. Renal cell carcinoma in tuberous sclerosis complex. Genes (Basel). 2021;12(10):1585.

4. Tapon N, Ito N, Dickson BJ, Treisman JE, Hariharan IK. The Drosophila tuberous sclerosis complex gene homologs restrict cell growth and cell proliferation. Cell. 2001;105(3):345–55.

5. Saxton RA, Sabatini DM. mTOR signaling in growth, metabolism, and disease. Cell. 2017;169(2):361–71.

6. Kim J, Kundu M, Viollet B, Guan KL. AMPK and mTOR regulate autophagy through direct phosphorylation of Ulk1. Nat Cell Biol. 2011;13(2):132–41.

7. Parkhitko A, Myachina F, Morrison TA, Hindi KM, Auricchio N, Karbowniczek M, et al. Tumorigenesis in tuberous sclerosis complex is autophagy and p62/sequestosome 1 (SQSTM1)-dependent. Proc Natl Acad Sci U S A. 2011;108(30):12455–60.

8. Rosset C, Jaeger MDC, Filippi-Chiela E, Reis LB, Sartor ITS, Oliveira Netto CB, et al. Primary cells derived from Tuberous Sclerosis Complex patients show autophagy alteration in the haploinsufficiency state. Genet Mol Biol. 2021;44(4):e20200475.

9. Knudson AG Jr. Mutation and cancer: Statistical study of retinoblastoma. Proc Natl Acad Sci U S A. 1971;68(4):820–3.

10. Alquezar C, Schoch KM, Geier EG, Ramos EM, Scrivo A, Li KH, et al. TSC1 loss increases risk for tauopathy by inducing tau acetylation and preventing tau clearance via chaperone-mediated autophagy. Sci Adv. 2021;7(45):eabg3897.

11. Franz DN, Belousova E, Sparagana S, Bebin EM, Frost MD, Kuperman R, et al. Long-term use of everolimus in patients with tuberous sclerosis complex: final results from the EXIST-1 study. PLoS One. 2016;11(6):e0158476.

12. Henske EP, Jóźwiak S, Kingswood JC, Sampson JR, Thiele EA. Tuberous sclerosis complex. Nat Rev Dis Primers. 2016;2:16035.

14. Klionsky DJ, Petroni G, Amaravadi RK, et al. Autophagy in major human diseases. EMBO J. 2021;40(19):e108863.

15. Tyburczy ME, Dies KA, Glass J, Camposano S, Chekaluk Y, Thorner AR, et al. Mosaic and intronic mutations in TSC1/TSC2 explain the majority of TSC patients with no mutation identified by conventional testing. PLoS Genet. 2015;11(11):e1005637.

16. Demidenko ZN, Zubova SG, Bukreeva EI, Pospelov VA, Pospelova TV, Blagosklonny MV. Rapamycin decelerates cellular senescence. Cell Cycle. 2009;8(12):1888–95.

17. Dowling RJ, Zakikhani M, Fantus IG, Pollak M, Sonenberg N. Metformin inhibits mammalian target of rapamycin-dependent translation initiation in breast cancer cells. Cancer Res. 2007;67(22):10804–12.

18. Meric-Bernstam F, Gonzalez-Angulo AM. Targeting the mTOR signaling network for cancer therapy. J Clin Oncol. 2009;27(13):2278–87.

19. Amin S, Mallick AA, Edwards H, Cortina-Borja M, Laugharne M, Likeman M, et al. The metformin in tuberous sclerosis (MiTS) study: A randomised double-blind placebo-controlled trial. EClinicalMedicine. 2021;32:100715.

20. Wu H, Ding J, Li S, Lin J, Jiang R, Lin C, et al. Metformin promotes the survival of random-pattern skin flaps by inducing autophagy via the AMPK-mTOR-TFEB signaling pathway. Int J Biol Sci. 2019;15(2):325–40.

21. Richards S, Aziz N, Bale S, Bick D, Das S, Gastier-Foster J, et al. Standards and guidelines for the interpretation of sequence variants: a joint consensus recommendation of the American College of Medical Genetics and Genomics and the Association for Molecular Pathology. Genet Med. 2015;17(5):405–24.

22. Haapaniemi E, Botla S, Persson J, Schmierer B, Taipale J. CRISPR-Cas9 genome editing induces a p53-mediated DNA damage response. Nat Med. 2018;24(7):927–30.

23. Seki A, Rutz S. Optimized RNP transfection for highly efficient CRISPR/Cas9-mediated gene knockout in primary T cells. J Exp Med. 2018;215(3):985–97.

24. Sherr CJ, DePinho RA. Cellular senescence: mitotic clock or culture shock? Cell. 2000;102(4):407–10.

25. Gorgoulis V, Adams PD, Alimonti A, Bennett DC, Bischof O, Bishop C, et al. Cellular senescence: defining a path forward. Cell. 2019;179(4):813–27.

