## Supplemental material for "Metformin modulates autophagy in heterozygous and CRISPR-edited *TSC2* primary fibroblasts"

**Supplementary table 1. Immunofluorescence quantification.**

| LC3-II mean (A.U) | c.724A>T het | c.724A>T CR | c.1008T>G het | c.1008T>G CR | c.4375C>T het | c.4375C>T CR |
|---|---|---|---|---|---|---|
| DMSO | 748088 | 913170 | 7372216 | 921588 | 1356313 | 1587059 |
| Rapamycin | 887560 | 1526087 | 918627 | 1487227 | 1042444 | 22239235 |
| Metformin | 954036 | 1418895 | 1147204 | 1392122 | 1643433 | 2926077 |
| HBSS | 1366486 | 1815913 | 1133797 | 1751182 | 2060003 | 2668270 |
| **p-s6k mean (A.U)** | | | | | | |
| DMSO | 621411 | 601190 | 582146 | 559677 | 1606479 | 1429240 |
| Rapamycin | 489071 | 274973 | 332109 | 249976 | 962404 | 594746 |
| Metformin | 431041 | 350555 | 256869 | 202777 | 1125363 | 849732 |
| HBSS | 180300 | 156945 | 180300 | 156945 | 474719 | 498225 |
| **N** | | | | | | |
| DMSO | 376 | 307 | 326 | 230 | 366 | 418 |
| Rapamycin | 439 | 263 | 246 | 284 | 395 | 461 |
| Metformin | 418 | 256 | 342 | 243 | 317 | 310 |
| HBSS | 482 | 236 | 303 | 243 | 469 | 263 |
