## Supplemental material for "Metformin modulates autophagy in heterozygous and CRISPR-edited *TSC2* primary fibroblasts"

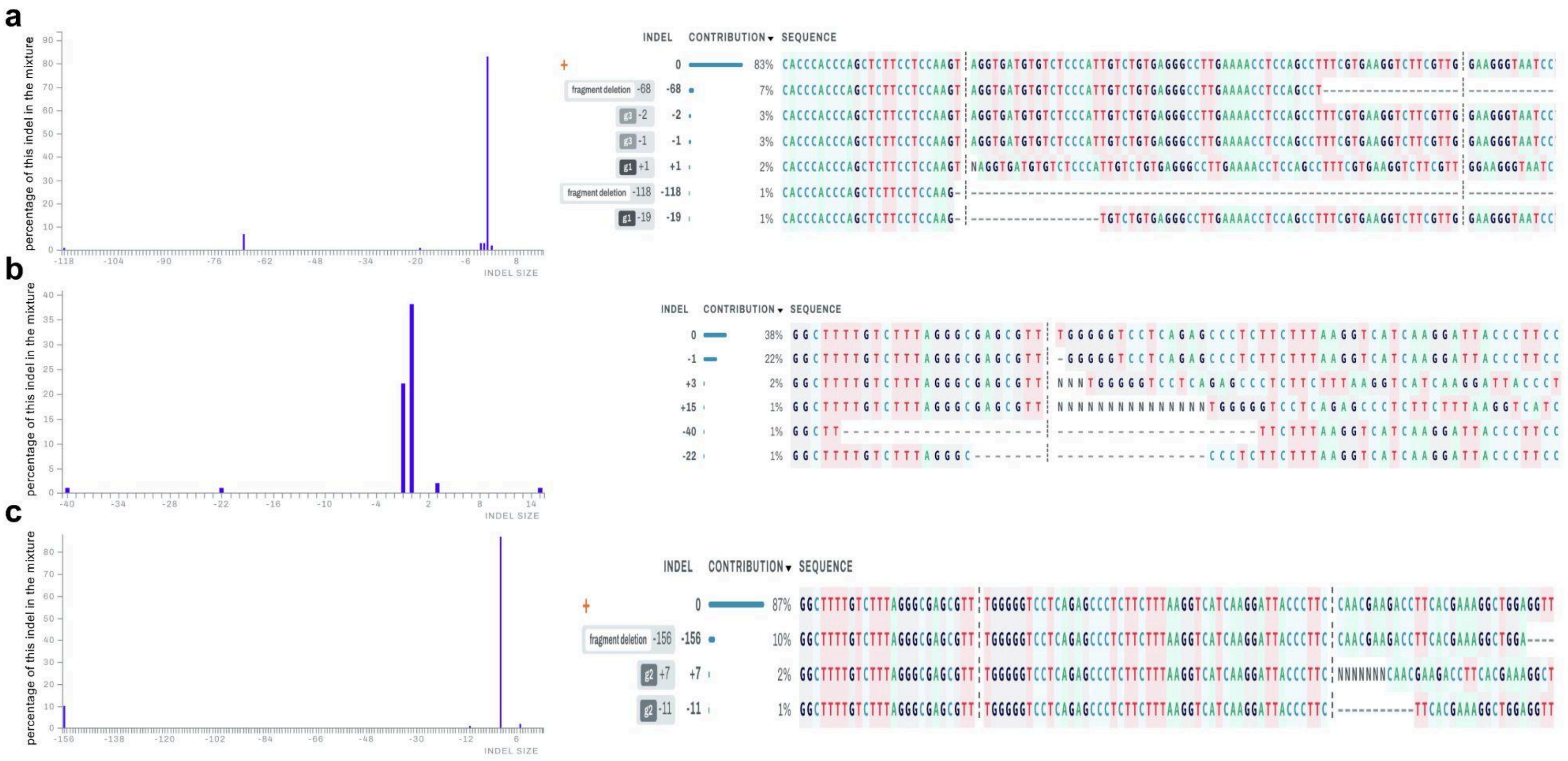


Supplementary Figure 1. Edited population for each cell line. Percentage of indels in the mixture and where each edition was made in the genome. 0= unedited population. g1,g2, g3 is the guide interference in the edition and fragment deletion is the consequence of multiple guides action in the same strand.

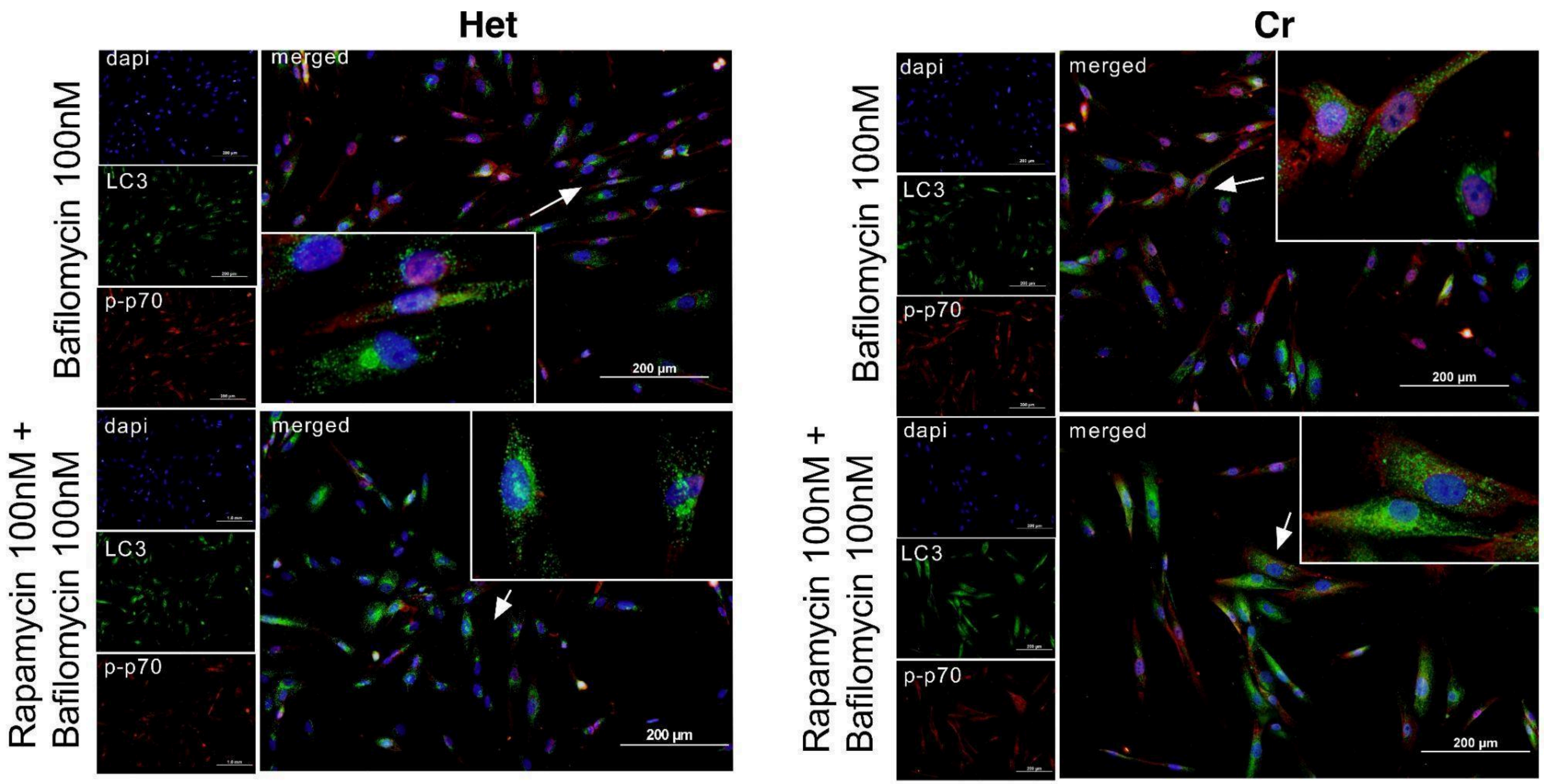


Supplementary Figure 2. Visual characterization of autophagic flux and mTORC1 activity by immunofluorescence. Representative fluorescence microscopy images of heterozygous (Het) and CRISPR-edited (Cr) TSC2 primary fibroblasts. Cells were treated with Bafilomycin A1 (100nM) alone or in combination with Rapamycin (100nM). Staining was performed for DAPI (blue), indicating nuclei; LC3 (green), p-p70S6K (red). White arrows and insets highlight the cells with accumulation of green puncta (LC3-II) within the cytoplasm.
